# The Genetic Diversity of Two Subpopulations of Shenxian Pigs as Revealed by Genome Resequencing

**DOI:** 10.1101/2021.01.25.428118

**Authors:** Liu Diao, Cao Hongzhan, Lu Chunlian, Li Shang, Jia Mengyu, Li Sai, Ren Liqin

**Affiliations:** College of Animal Science and Technology, Hebei Agricultural University, Baoding 071000, Hebei; Hebei Zhengnong Animal Husbandry Co., Ltd., Xinji 052360, Hebei; Fishery Technology Extension Station, Wanquan District, Zhangjiakou City, Zhangjiakou, Hebei 076250

**Keywords:** Genome resequencing, Shenxian pig, genetic diversity, analysis

## Abstract

Shenxian pigs can be divided into two main strains from their shape and appearance: Huangguazui and Wuhuatou. There are significant differences in the phenotypic characters between the two subpopulations. The Shenxian pig, as the only local pig breed listed in China Pig Breeds in Hebei Province, has the excellent traits of Chinese local pigs. In order to explore the genetic diversity and genetic distance of the different subpopulations of Shenxian pig, as well as understand their evolutionary process, whole genome resequencing and genetic structure analyses were performed for the two sub-population types of Shenxian pigs.The results showed that a total of 14,509,223 SNP sites were detected in the Huangguazui type and 13,660,201 SNP sites were detected in the Wuhuatou type. This study’s principal component analysis results showed that the genetic differentiation of the Shenxian pig population was serious, and there was obvious stratification in the population. The phylogenetic tree analysis results indicated that there was a certain genetic distance between the Huangguazui and Wuhuatou types.

## INTRODUCTION

Shenxian pig is the only outstanding local pig breed listed in the “Chinese Pig Breeds” in Hebei Province. Compared with other imported pig breeds, Shenxian pigs have an earlier sexual maturity, short estrus interval, and a large number of litters. They have good research significance. There are two main strains of Shenxian pigs: Cucumber beak and Wuhuatou. “Cucumber Mouth”: The mouth is long and pointed at the end, hence the name. The nose is straight and flat, with few hairs, shallow wrinkles on the forehead, slender neck, narrow forequarters, long and concave back and waist, slightly drooping abdomen, inclined buttocks, weak hind limbs, and thick tail. “Five-flowered head”: The head is rough, the hair is thick, the forehead is wide and wrinkled, and it gets its name. Based on the whole-genome resequencing technology, the genetic structure of the two sub-populations of Shenxian pigs was discussed, and the evolutionary mechanism between the two populations was revealed at the molecular level, and all genetic information in the whole genome was accurately obtained, and the two populations were discovered to the greatest extent. Genetic variation(Bi et al.2020).

## MATERIALS AND METHODS

### Materials

The pig farm in Shenxian County was selected from Hebei Zhengnong Animal Husbandry Co., Ltd. and selected healthy, weight, and age-similar cucumber beaks and 5 pigs with five-flower head each. Cucumber beaks: B1-1, B1-2, B1-3, B1 -4, B1-5, five flower head: B2-1, B2-2, B2-3, B2-4, B2-5. Use ear tag tongs to collect ear tissue, shear pig ear tissue and disinfect with 75 percent alcohol, take about 100mg sample, store the sample in an EP tube containing 95 percent alcohol, transport it back to the laboratory with an ice pack, and store it in a refrigerator at -20°C It is reserved for use in sampling, transportation and storage to ensure that the samples are not contaminated for subsequent extraction of genomic DNA.

### Whole genome resequencing

Whole genome resequencing is process used to sequence the genomes of different individuals of species with known genome sequences, and then analyze the differences of individuals or groups on that basis(Ley et al.2008). It has been found that based on whole genome resequencing technology, researchers have been able to quickly carry out resource surveys and screening processes in order to determine large numbers of genetic variations, and realize genetic evolution analyses and predictions of important candidate genes. In the current study, the two subpopulations of Shenxian pigs were re-sequenced for the purpose of obtaining the genomic information. A large number of high accuracy SNPs, InDel, and other variation information were obtained by comparing the results with the reference genome. Subsequently, the variation information was successfully detected, annotated, and counted.

### Detection methods based on next generation sequencing technology

At present, the second-generation sequencing technology is widely used(Levy et al.2016;Slatko et al.2018).The development of the next generation sequencing (NGS) technology has resulted in revolutionary changes in the detections of genetic variations. Researchers can now comprehensively and accurately detect various types and the sizes of genetic variations from the genomic level(Mardis.2008) For the detection of genetic variations based on the next generation sequencing technology, the first step is to compare the reading segments of thousands of sequences to the corresponding positions on the reference genome. Then, the possible variation information can be inferred using simple mathematical models(Andersson.2009;Mckenna et al.2010;Montgomery et al.2013). These methods take the number of reading segments supporting the variations and reference sequences; quality of the reading segment comparison; environmental conditions of the genome sequences; and the possible noise (such as sequencing errors) as prior information. Then, the variation types and genotypes at certain positions can be determined according to naive Bayesian theory, and the genotype with the highest posterior probability can be successfully obtained(Montgomery et al.2013;Durbin et al.2008). Such methods have been found to be effective for SNP and small InDel detections(Depristo et al.2011;Albers et al.2011).

### Data processing

This test sample uses the whole genome resequencing data (sequencing depth 10X) of 10 Shenxian pigs (5 cucumber-mouthed pigs and 5 Wuhuatou pigs). The output of the genome data comparison is the sam file. Since the sorting method of the sam file cannot be used for subsequent analysis, the Samtools software(Li et al.2009)is used to convert the sam file into a file in chromosome sorting format, that is, the bam format. Chromosome rearrangement is to arrange the entries of the same chromosome in the file in ascending order, and merge the two files of the same sample.

SNP detection and annotation: Use GATK and VarScan software(Koboldt et al.2009)to perform SNP detection at the same time, so as to ensure that the obtained SNP site information will not be affected by the deviation of base misalignment caused by InDel mutation. GATK detection SNP code: java -jar GenomeAnalysisTK.jar glm SNP -R ref.fa -T UnifiedGenotyper -I test.sorted.repeatmark.bam -o test.raw.vcf. In order to ensure the accuracy of SNP information, strict testing conditions are set when detecting SNP site information: <1>The minimum number of end reads greater or equal to 4; <2>The minimum quality value Q20 greater or equal to 90; <3>Minimum coverage greater or equal to 6; <4>P value Less than or equal to 0.01. InDel detection and annotation: The same use of GATK and VarScan software for InDel detection can eliminate errors caused by SNP variation. GATK detection InDel code: java -jar GenomeAnalysisTK.jar glm InDel -R ref.fa -T UnifiedGenotyper -I test.sorted.repeatmark.bam -o test.raw.vcf. The detection conditions are consistent with SNP. The detected SNP and InDel mutation sites also need quality control, remove untrusted data sites, correct the quality value, and output the file in VCF format after correction. Use R language(Gao et al.2014;Sudhaka.2018)and ANNOVAR software (Kai et al.2014)to perform mutation information Sort and comment.

## RESULTS AND DISCUSSION

### Results

BWA software (Version: 0.7.15-r1140) was used in this research investigation to compare the clean sequences of the two subpopulations of Shenxian pigs after quality control with the latest version of reference genome of pigs was implemented. The results are detailed in Table 2-1. It was found that on average, more than 86 percent of the sequencing data could be compared to the reference genome.

There are a large number of SNPs, and there may be one SNP variant site every 500-1000bp(Group et al.2001).Count the gene region and type of SNP to get Table 1.A total of 14509223 SNPs were detected in Huangguazui pigs (B1), of which there were 88176.4 (0.61%) and 101948.2 (0.70%) in the upstream and downstream 1kb region of the gene, and 7865381 (54.21%) in the intergenic region. There are an average of 99217.4 (0.68%) and 6151792 (42.40%) of exons and introns. A total of 13660201 SNPs were detected in Wuhuatou pig (B2), of which 84855.2 (0.62%) and 97797.2 (0.72%) were found in the upstream 1kb region and the downstream 1kb region of the gene, and the intergenic region contained 7407500 (54.23%). There are 96530.8 (0.71%) and 5782023 (42.33%) of exons and introns respectively.

**Table 1.** Statistical table of gene region and type of SNP.

| Sample-name | Downstream | Exonic | Intergenic | Intronic | Upstream | Total<sup>a</sup> |
| --- | --- | --- | --- | --- | --- | --- |
| Huangguazui(B1) | 101948 0.70% | 99217 0.68% | 7865381 54.21% | 6151792 42.40% | 88176 0.61% | 14509223. |
| Wuhuatou(B2) | 97797 0.72% | 96530 0.71% | 7407500 54.23% | 5782023 42.33% | 84855 0.62% | 13660201. |

<sup>a</sup> There is little difference in the number of SNPs contained in the cucumber mouth and five-flower head pig gene regions. Introns and intergenic regions contain the most SNPs, indicating that the two regions have the most variation.

In the current study, the SNP was identified using GATK software, and the SNP data were subsequently analyzed. The original data contained a total of ten samples and 31,658,350 SNP sites. Effective data was obtained after a secondary quality control of the original data, with ten samples and 23,088,823 SNP sites remaining. The total samples contained the individuals of the two strains of Shenxian pigs.A total of 580 high-quality polymorphic SNPs were obtained by screening 23,088,823 SNP sites. The marker covered eleven chromosomes, and its PIC index ranged from 0.1 to 0.529, with an average value of 0.305. There were 119 highly polymorphic SNPs and 191 moderately polymorphic SNPs identified.In this study, the quantity and density distributions of the SNPs on chromosomes were further analyzed. The results are shown in Figure 1. The distributions of the quantities and densities of the SNPs on the chromosomes were observed to not be uniform. For example, Chromosome 15 was short in length but had the largest number of SNPs, which was far more than the other examined chromosomes. These findings indicated that the densities of the SNPs on that particular chromosome were the highest. Therefore, it was considered that there was a relatively high density of variation on Chromosome 15.

**Figure 1.**
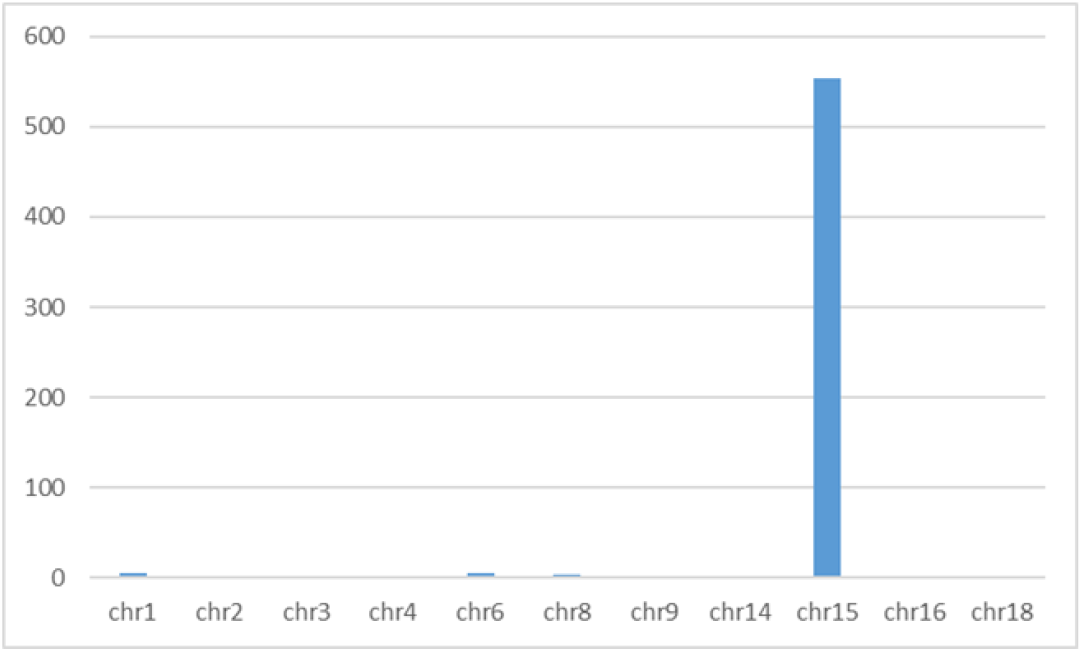
The abscissa represents the chromosome number, and the ordinate represents the number of SNPs.The distributions of the quantities and densities of the SNPs on the chromosomes were observed to not be uniform. For example, Chromosome 15 was short in length but had the largest number of SNPs, which was far more than the other examined chromosomes. These findings indicated that the densities of the SNPs on that particular chromosome were the highest. Therefore, it was considered that there was a relatively high density of variation on Chromosome 15.

In order to infer the genetic relationship between the Huangguazui and Wuhuatou pig subpopulations, a genetic distance matrix of the IBS was constructed using Plink (V1.9) software(Purcell et al.2007). Then, based on the results, a phylogenetic tree was constructed using a neighbor joining method (NJ), as detailed in Figure 2. The analysis results showed that the Huangguazui and Wuhuatou subpopulations had originated from the same ancestor and belonged to the same variety. However, they could clearly be divided into two distinct groups, which indicated that there were indeed differences in the genetic backgrounds of the Huangguazui and Wuhuatou pigs. It was confirmed that there was a certain distance between the two strains in their genetic relationship.It was also determined in the phylogenetic tree analyses of the Shenxian pigs, large white pigs representative of introduced pigs, and Meishan pigs representative of local south China pigs, that the three populations had originated from the same species, and there was a distant genetic relationship between the Shenxian pigs and the other two examined populations.

**Figure 2.**
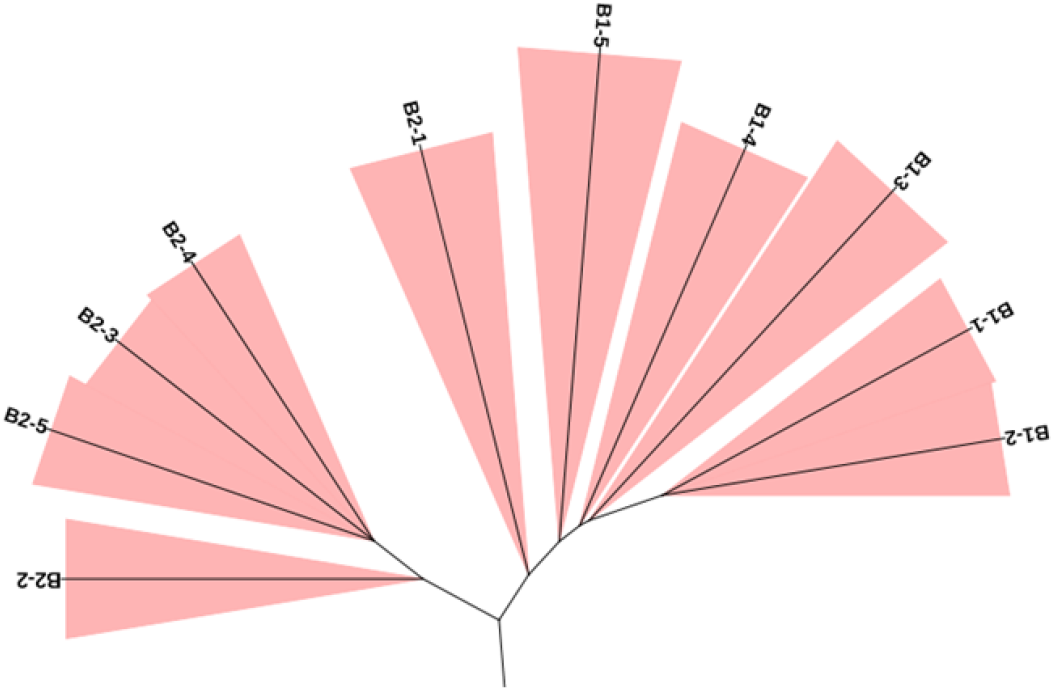
The analysis results showed that the Huangguazui and Wuhuatou subpopulations had originated from the same ancestor and belonged to the same variety. However, they could clearly be divided into two distinct groups, which indicated that there were indeed differences in the genetic backgrounds of the Huangguazui and Wuhuatou pigs. It was confirmed that there was a certain distance between the two strains in their genetic relationship.

The PCA analysis of the effective data obtained after quality controls were implemented were conducted using Plink software, and then the PCA analysis results were visualized using R software. In order to reveal the population specificity of the Huangguazui and Wuhuatou pigs, the SNP information of the two populations was analyzed using a principal component analysis (PCA) method. The top three eigenvectors were selected for the analysis, and the analysis results were visualized by R language, as shown in Fig. 2-3. The analysis results of the first and second eigenvectors are detailed in Figure 3. It can be seen in the figure that the first eigenvector had accurately distinguished the Huangguazui and Wuhuatou into two strains, which confirmed that the differentiation among the Shenxian pig population was serious. The second eigenvector revealed that both the Huangguazui and Wuhuatou subpopulations were stratified within the total population. The analysis results of the first and third eigenvectors showed that the stratification in Huangguazui population was the most obvious (Figure 4). The second and third eigenvectors showed (Figure 5) that the degrees of stratification in the Wuhuatou strain were less than those of the Huangguazui strain.

**Figure 3.**
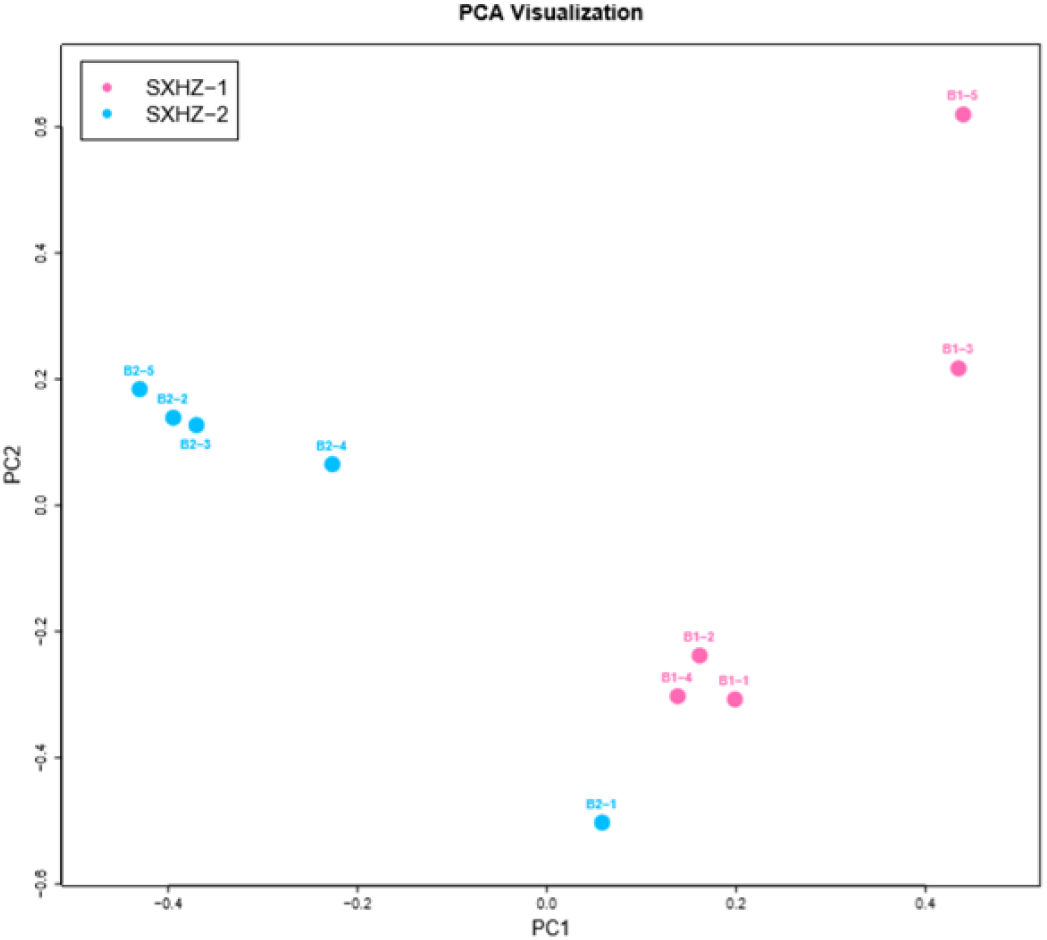
The abscissa represents the first vector, and the ordinate represents the second vector; pink represents cucumber-mouthed pigs, blue represents five-flowered pigs.

**Figure 4.**
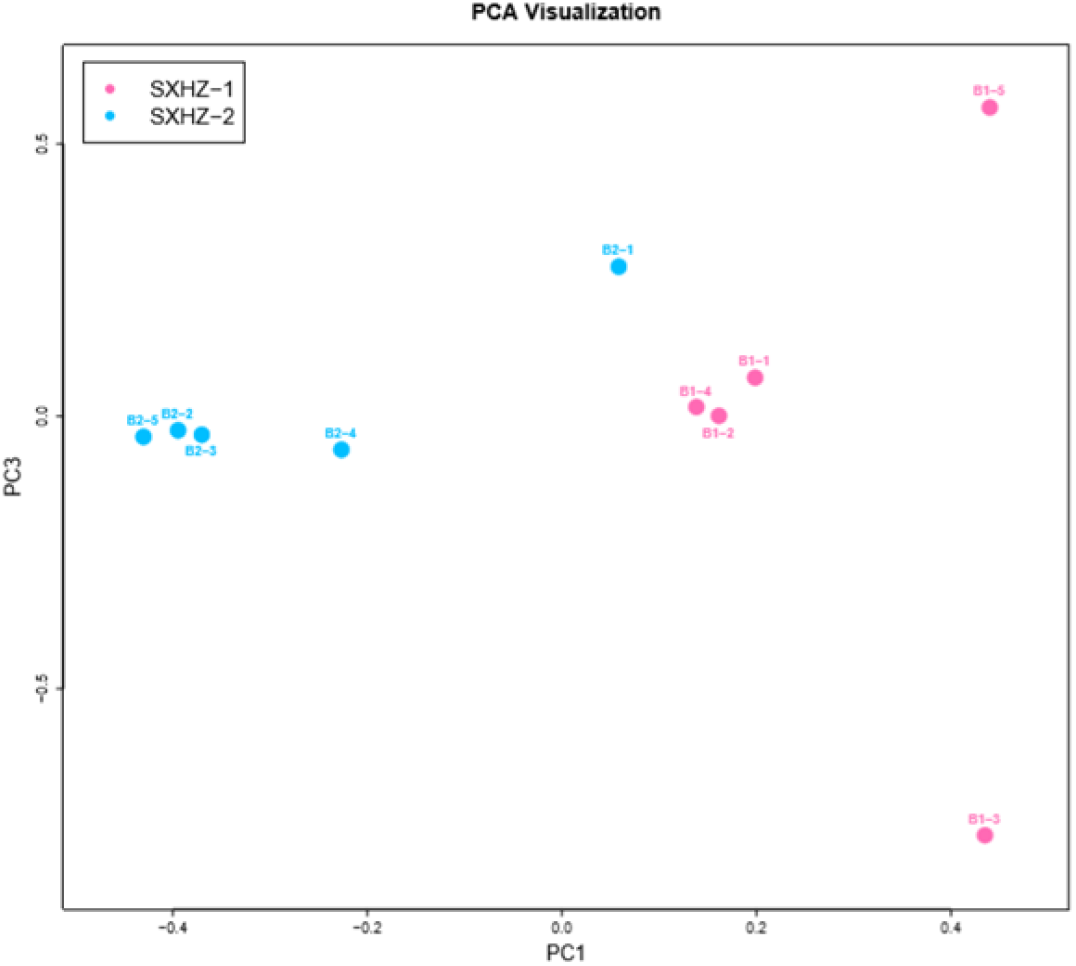
The abscissa represents the first vector, and the ordinate represents the third vector; pink represents cucumber-mouthed pigs, blue represents five-flowered pigs.

**Figure 5.**
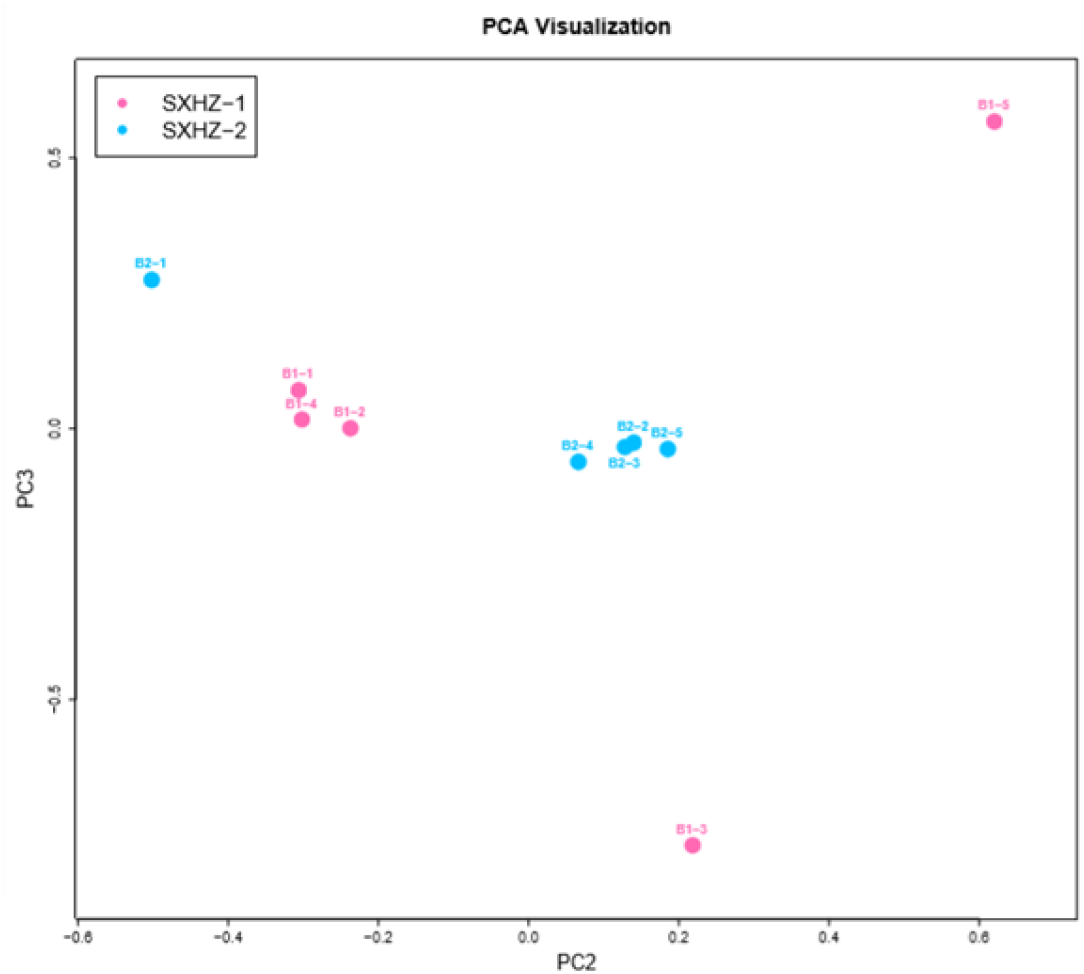
The abscissa represents the second vector, and the ordinate represents the third vector; pink represents cucumber-mouthed pigs, blue represents five-flowered pigs.

## Discussion

For DNA extraction and detection, the complete DNA in the agarose gel should be a band. At times, there will be three bands observed, which may have been formed by DNA degradation. Generally speaking, the foremost band is of supercoiled configuration; middle band is linear; and the last band is of single-chain defective configuration(Martina et al.2020). If mechanical damages occur during the extraction process, the DNA can become degraded, which may be presented as a diffuse state on the gel.

SNP mainly refers to the DNA sequence polymorphism caused by a single nucleotide variation at the genomic level. In this study, the SNP information obtained via statistical variation detection showed that the quantity and density distributions of the SNPs in the two subpopulations were uneven, and they were mainly concentrated on Chromosome 15. There have been many studies conducted regarding the QTL of meat quality traits on Chromosome 15(Liu.2015;Jia.2004), which may be related to the excellent meat quality traits of the Shenxian pig species(Hu et al.2015). Previous studies have successfully mapped the QTLs of the meat quality traits and acquired linked SNPs for labelling(Yan et al.2017;Li et al.2010;Science.2016b). However, the current SNP chip technology is not adequately developed for mapping the quantitative traits in pigs(Chen et al.2011). Therefore, the SNP and InDel information on Chromosome 15 will be detected, annotated, and analyzed in follow-up experimental research.

Furthermore, functional gene annotations will also be carried out in order to explore the differential genes, along with the quality traits regulated by the differential genes, in the two subpopulations of Shenxian pigs. Another area to be explored in the future will be the main traits controlled by Chromosome 15 in order to provide a research basis for the follow-up breeding of Shenxian pigs.

The whole genome population structure analysis processes are utilized to re-sequence different geographic locations and populations, as well as to obtain the SNP and other information after comparisons are made with the reference genome. Subsequently, the principal components and population structures among populations are analyzed, and appropriate phylogenetic trees are constructed. In the present study, based on the analysis results of the population structures of the examined species, the differences in the members of the examined species could be more clearly understood. It was found that the genome-wide population structure analysis results had inferred the characteristics, population structure, and development history of the varieties from the genome. The obvious stratification and severe phenotypic differentiation observed in the population structures of the Shenxian pigs indicated that the phenotypic traits of Huangguazui and Wuhuatou may have been less affected by artificial selection processes.

